# Perceptual decisions are biased toward relevant prior choices

**DOI:** 10.1101/858324

**Authors:** Helen Feigin, Shira Baror, Moshe Bar, Adam Zaidel

## Abstract

Perceptual decisions are biased by recent perceptual history – a phenomenon termed ‘serial-dependence.’ Using a visual location discrimination task, we investigated what aspects of perceptual decisions lead to serial dependence, and disambiguated the influences of low-level sensory information, prior choices and motor actions on subsequent perceptual decisions. Following several biased (prior) location discriminations, subsequent (test) discriminations were biased toward the prior choices, even when reported via different motor actions, and when prior and test stimuli differed in color. By contrast, biased discriminations about an irrelevant stimulus feature did not substantially influence subsequent location discriminations. Additionally, biased stimulus locations, when color was discriminated, no longer substantially influenced subsequent location decisions. Hence, the degree of relevance between prior and subsequent perceptual decisions is a key factor for serial-dependence. This suggests that serial-dependence reflects a high-level mechanism by which the brain predicts and interprets incoming sensory information in accordance with relevant prior choices.

## INTRODUCTION

Perceptual decisions are not based solely on current sensory information. Rather, they are substantially influenced by other factors, such as context and prior experience (Kersten et al., 2004; Bar, 2004; Panichello et al., 2013; Clark, 2013; de Lange et al., 2018). Notably, our perceptual decisions are biased by preceding perceptual events – a phenomenon, termed ‘serial-dependence’, that has garnered considerable recent interest (Fischer and Whitney, 2014; Liberman et al., 2014; Taubert et al., 2016; Alexi et al., 2018; Liberman and Manassi, 2018). Serial-dependence can help or impair performance, depending on circumstances. For example, when a discriminated feature is distributed independently over time (or across trials, as is often the case in laboratory experiments) preceding information has no bearing on current stimuli, and therefore serial-dependence adds noise and degrades performance (Abrahamyan et al., 2016). By contrast, events in the real world are generally not independent in that they follow prototypical regularities in time and space. Hence, serial-dependence might benefit everyday (ecological) performance by supporting perceptual stability over time (Fischer and Whitney, 2014; Liberman et al., 2016; Manassi et al., 2017), and by improving speed and efficiency of perceptual decisions (Stefanics et al., 2010; Burr and Cicchini, 2014; Cicchini et al., 2014). Overall, serial-dependence, like predictions more generally, reflects a trade-off between efficiency (relying on biased expectations) and accuracy (relying on instantaneous sensory information).

The specific conditions that give rise to serial-dependence are not fully understood. Many factors may trigger it, including sensory inputs (Fischer and Whitney, 2014; Cicchini et al., 2017), motor responses (de Lange et al., 2013; Pape and Siegel, 2016), expectations (Chalk et al., 2010; Kok et al., 2013) and spatial attention (Fischer and Whitney, 2014; Morales et al., 2015). It has even been proposed that prior choices, in and of themselves, can elicit serial-dependence (Kaneko and Sakai, 2015; Talluri et al., 2018). However, the nature of the elements that underlie serial-dependence and their relative contribution to this phenomenon are not fully known.

If serial-dependence is harnessed by the brain to aid function and efficiency, then it should only be used when preceding perceptual events are *relevant* to the impending perceptual decision. By *relevant*, we imply that the observer presumes that the series of observations are statistically related, and therefore preceding events can, on average, aid subsequent decisions. When irrelevant, preceding events should have little or no effects on subsequent perceptual decisions. Accordingly, we hypothesized that relevance between sequential perceptual events may be a key factor that determines whether or not serial-dependence would occur. In support of this idea Suárez-Pinilla et al (2018) recently found that estimates of a stimulus feature (variance) were biased only when that same feature, but not when a different feature (mean), was discriminated. By contrast, Fornaciai and Park (2018a, 2018b) did find serial-dependence when participants were told that the prior stimuli were irrelevant, and to ignore them. Therefore, the effects of relevance on serial-dependence require further investigation. Moreover, for a complete understanding, specific contributions to serial-dependence of low-level sensory information, motor actions, and choices (when relevant or irrelevant) require disambiguation.

For this purpose, we devised a novel paradigm of visual location discrimination that induces short-term biases over several (prior) stimuli, and tested their effects in an interleaved and balanced manner. To examine how the relevance of perceptual decisions affects serial-dependence, we manipulated specific features of the prior decisions including stimulus characteristics, which feature was discriminated, and the motor action used to report choices. We refer to the process of discriminating a stimulus feature as a *perceptual decision*, and the result of the process as the *choice*, which may (or many not) be reported via a motor response.

We found that recent perceptual decisions elicit serial-dependence only when relevant. This was seen even when the prior choices were reported via different motor actions, and when the prior stimuli had different sensory attributes (color) from the test stimuli. As long as the same feature (location) was being discriminated in both prior and test stimuli, serial-dependence was observed. By contrast, irrelevant prior decisions (about stimulus color) did not bias subsequent location discriminations, even when the prior motor actions or prior stimulus locations (which were not discriminated) were biased. Hence, relevance (i.e., decisions about the same feature of interest) is a key element for serial-dependence.

## METHODS

### Participants

A total of 114 participants took part in the study (68 females; mean age ± SD: 24 ± 3.54, range: 18-37 years). All participants were healthy and had normal or corrected-to-normal vision. This study was approved by the Bar-Ilan University, Gonda Brain Research Center Ethics Committee. All participants signed informed consent and received monitory compensation for their participation. The data gathered were pre-screened to confirm task understanding and cooperation (see *Statistical analysis* section below for details). This excluded 11 participants’ data, leaving 103 for further analysis. Additional participant details, pertaining to specific experimental conditions, are presented below in the section *Experimental conditions*. The sample size of our study is in line with previous studies exploring serial-dependence in perceptual decisions (e.g. Fornaciai & Park, 2018)

### Experimental design

Stimuli were generated and presented using PsychoPy software (Peirce, 2007) on an LCD monitor with a resolution of 1920×1200 pixels and a refresh rate of 60 Hz. Experiments were run in a dark, quiet room. Participants were seated in front of the monitor, with their head positioned 45 cm from the center of the screen and supported by a chin rest. Stimuli comprised uniformly filled circles (6.36° visual angle) presented around the center of the screen on a dark gray background. Circle location was varied in the horizontal dimension only. Participants performed a two-alternative forced choice (2AFC) task in which they discriminated the location of the circle presented (left or right relative to the screen’s center) by pressing the corresponding arrow key on a standard computer keyboard. Participants were instructed to make their choice as rapidly and accurately as possible. They were informed that task difficulty varies with circle proximity to the screen center and instructed to make their best guess when in doubt.

Each trial comprised a series of several ‘prior’ circle stimuli, followed by a single ‘test’ circle stimulus (each discriminated individually, by sequence). The prior stimuli in a given trial were drawn from a specific distribution that was biased (either leftward or rightward) or unbiased. The ‘test’ stimulus was unbiased. This novel paradigm allowed us to examine the rapid effects of (short-term) biases, in a controlled and interleaved manner. Figure 1 shows the event sequence of a single trial. A fixation cross appeared for 500 ms at the beginning of each trial. This marked the center of the screen, and was presented at the beginning of each trial in order to reduce possible carry-over effects from the previous trial. The fixation cross was not presented again during the rest of the trial in order not to interfere with the priors’ effect being measured. Each circle (prior/test) was presented for 250 ms, and was preceded by a 250 ms delay (blank screen) such that the next circle appeared 250 ms after the participant’s previous response.

**Figure 1.**
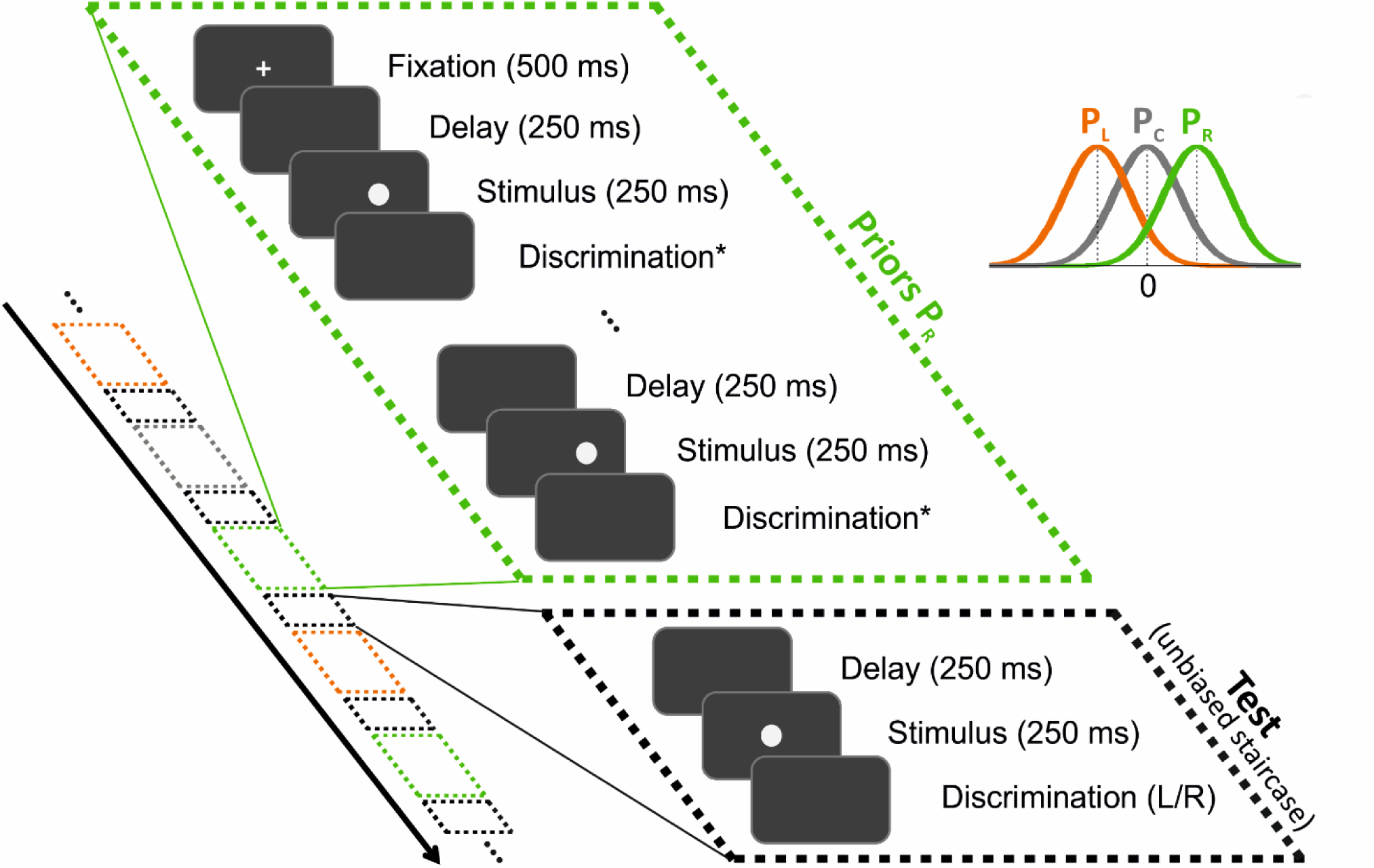
Event sequence within and across trials. Each trial began with a fixation point (at the screen center) followed by a series of discriminations of ‘prior’ stimuli (for example, priors biased to the right, green dashed boxes), and then discrimination of a single ‘test’ stimulus (black dashed boxes). The black arrow marks the timecourse of the experiment, which interleaved trials of different prior types pseudorandomly (right, left and center, marked by green, orange and gray boxes, respectively), each ending with a test stimulus (black box). The stimuli were circles on a dark gray screen, and were discriminated individually (in order). The discrimination task for the test circles was about location: was the circle to the left or to the right of the sceen center? For the prior circles, the discrimination task (*) depended on the experimental condition. It was either location discrimination (like the test task) or color discrimination (unlike the test task). All discriminations required choosing one from two alternatives: right/left or green/purple (for location and color discrimination, respectively). Prior circle locations were drawn from a normal distribution and were either biased to the left or right of the screen center (orange and green probability distributions, P_L_ and P_R_, respectively) or unbiased (center distribution P_C_, in gray). Test circle locations followed an unbiased starcaise procedure.

We defined the locations of the stimuli (circle center) in the horizontal plane (x) by its visual angle relative to the screen center (x = 0°), such that positive and negative values represent locations positioned to the right and to the left of the screen center, respectively. Locations on the vertical plane were fixed (screen centered). Prior circles had randomly distributed locations, which were either (on average) biased to the left (mean ± SD: −3.82 ± 3.82°) or to the right (mean ± SD: 3.82 ± 3.82°) of the screen center, or unbiased (mean ± SD: 0 ± 3.82°). The prior “type” was defined by its bias: ‘left’, ‘right’ and ‘center’, respectively. With this definition, biased priors could still lie to the contralateral side, though with lower probability (∼16%).

Test circle locations were balanced to the right/left (the location sign was randomly selected on each trial), with magnitude (absolute distance from 0°) set using a staircase procedure (Cornsweet, 1962). This began at a location magnitude of |x| = 6.36° and followed a “one-up, two-down” staircase rule. Namely, location magnitude |x| increased (i.e., became easier) after a single incorrect response, and decreased (i.e., became harder) after two successive correct responses. This staircase rule converges to 70.7% correct performance (Leek, 2001). Staircase step size followed logarithmic increments, such that location magnitude |x| was multiplied by 0.6 to increase task difficulty and divided by 0.6 to decrease task difficulty. Location magnitude was limited to a maximum of |x| = 25.46°, but this was never reached in practice. Separate staircase procedures were run for each prior type, each comprising 100 trials that were randomly interleaved. Thus, testing all three prior types (right, left and center) comprised 300 trials, and took ∼30 minutes, and testing only right/left priors comprised 200 trials and took ∼20 minutes. A break was automatically given after every 100 trials, and the experiment was resumed by the participant, when ready.

### Experimental conditions

Six experimental conditions were tested in order to differentiate the effects of different factors on serial-dependence. In four of the six conditions (conditions #1-4), the same stimulus feature (location) was discriminated for both prior and test circles. Hence, the prior decisions were considered relevant to the subsequent test circle perceptual decision. By contrast, in the other two conditions (conditions #5-6), a different feature (color) was discriminated for the prior circles, while location was still discriminated for the test circles. Therefore, the prior decisions were irrelevant to the subsequent test circle’s perceptual decision. The results of each such condition were analyzed separately. These six conditions are described here below in detail (and summarized in Table 1):

**Table 1.**
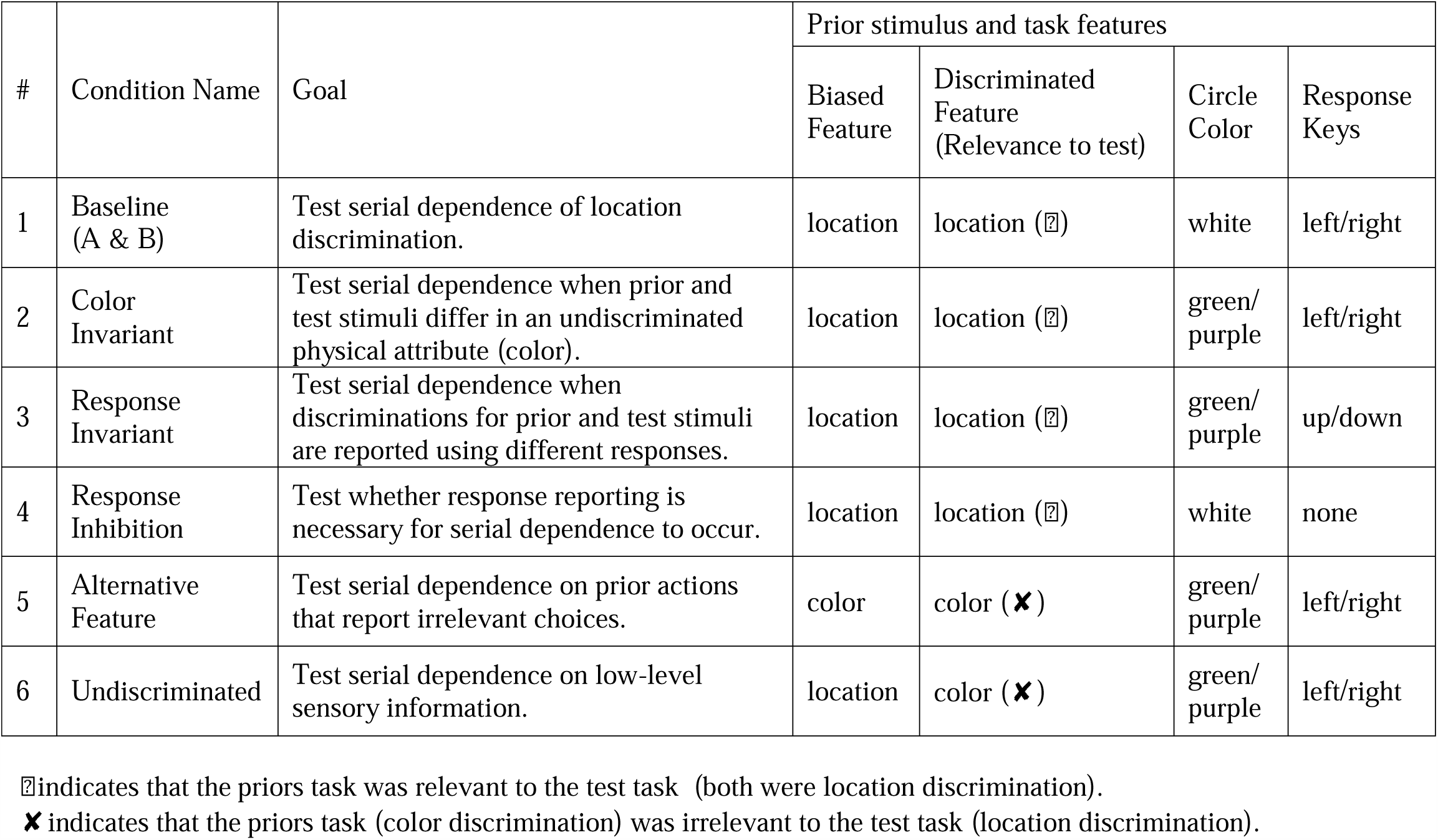
Experimental conditions.

### Condition #1: “Baseline”

In the baseline condition, the prior and test circle stimuli were indistinguishable (white circles). Participants were asked to perform the same task regarding all circles - to discriminate the location of each circle. We tested two variants for this condition: in *Baseline A.*, a fixed number of priors (*Q* = 5) preceded the test circle (the parameter *Q* reflects the number of priors on a given trial, and *N* the number of trials, which is also the number of test stimuli). In *Baseline B.*, the number of prior circles in each trial was random (*Q* was selected, per trial, according to a discretized Gaussian probability distribution with mean ± SD = 5 ± 2, limited within the range 1-9).We tested all three prior types (right/center/left) in *Baseline A*, but only the two biased prior types (right/left) for the rest of the experimental conditions (*Baseline B* and conditions #2-6), with 100 trials per prior type (i.e., *N* = 300 for Condition #1 *Baseline A* and *N* = 200 for the rest of the conditions). *Baseline A* and *B* were performed by two groups of 20 and 19 participants, respectively. In each variant of this condition, five participants were excluded from the analysis due to poor goodness-of-fit (see *Statistical analysis* section for details regarding the exclusion criterion). The group that took part in *Baseline B* also performed Condition #4 (*Response Inhibition*) during the same experimental session, as a separate block, with block order counterbalanced between participants.

### Condition #2: “Color Invariant”

In this condition, the same task (location discrimination) was tested for both prior and test stimuli, but the prior circles differed in color. The rationale for this condition was to test whether changing a minor (task-irrelevant) stimulus feature would affect serial-dependence. Also, since color was used in the following conditions to indicate a different task, this condition was required in order to rule out any differences due to color per se. Participants were instructed to discriminate the location of circles, irrespective of their color. This condition was essentially identical to the *Baseline B*, except that prior circles were colored (either green or purple) while test circles remained white. All stimuli (priors and test) had equal brightness. The color was randomly chosen for each prior stimulus (50% green and 50% purple), such that the color parameter was unbiased across trials. Twenty-four participants performed this condition; three participants were excluded from the analysis due to poor goodness-of-fit.

### Condition #3: “Response Invariant”

In this condition, the same task (location discrimination) was tested for both prior and test circles, but the choices for the prior and test circles were reported using different sets of keys. Like in condition #2 (*Color Invariant*) prior circles were colored (50% green and 50% purple) and the test circles were white. Participants were instructed to report right and left choices using the ‘up’ and ‘down’ arrow keys (counterbalanced) for colored circles, but using the ‘right’ and ‘left’ arrow keys for white circles. This condition allowed us to test whether serial-dependence ensues even when the actions for reporting choices are dissociated, but the high-level relevance of prior choices is maintained. Twenty-six participants performed this condition; four were excluded from the analysis due to poor goodness-of-fit. Twenty-five of the participants also performed Condition #5 (*Alternative Feature*) in a separate session that took place on a different day (with task order counterbalanced between participants), and 15 agreed to return also for a third session to perform Condition #6 (*Undiscriminated*).

### Condition #4: “Response Inhibition”

Here, prior and test stimuli were the same (white circles). However, participants were instructed to respond only to the last (i.e. the test) circle in each trial. The number of prior circles followed a random distribution (range 1-9, like *Baseline B*, and all the other conditions except for *Baseline A*). Therefore, the last circle could only be identified in retrospect (when it was not followed by another circle). Hence, the participants needed to discriminate all the circles, but suppress their responses to the prior circles, and respond only to the test circles. This condition allowed us to probe whether choices with unexecuted actions would still elicit serial-dependence. This condition was performed by the same participants from Condition #1 (*Baseline B*) in a counterbalanced order. Two participants were excluded from the analysis of this condition due to poor goodness-of-fit.

### Condition #5: “Alternative Feature”

The aim of this condition was to test whether irrelevant choices (i.e., relating to prior stimulus features other than location) with biased motor responses would elicit serial-dependence. Here, participants were instructed to report the color of the circle when it was green or purple (prior circles), but to discriminate location for the white (test) circles. Success in the color task was confirmed by a high percentage of correct responses (mean ± SD = 96.21 ± 2.46%). The prior circle colors in a given trial were biased, with an 84% probability for one color (16% for the other). Trials with prior circles biased green or purple were randomly interleaved in a block. The 84/16 percentage bias was chosen to match the ratio of right/left biased prior locations from the other conditions. Each color was coupled to one of the response keys (right/left arrow), counterbalanced across participants. Participants were instructed to also report the location of the white circle using the same right/left arrow keys. The locations of the prior circles were unbiased and followed the ‘center’ prior type distribution. Thus, the responses to prior stimuli were biased by color choices - irrelevant to the subsequent location discrimination for the test circle. This procedure therefore biased participants’ choices and motor actions in a manner which was irrelevant to the subsequent location discrimination.

This condition was performed by 25 participants who also performed Condition #3 (*Response Invariant*) in a different session (counterbalanced). Three participants were excluded due to poor goodness-of-fit and another two because of poor performance on the priors’ task (both below 50% correct, substantially lower than the rest of the participants, who all had above 89% correct).

### Condition #6: “Undiscriminated”

The aim of this condition was to test whether biased locations would still elicit serial-dependence when not actively discriminated. This was done by instructing the participants to report the color for the colored (prior) circles, and location for the white (test) circles. The colors of the prior circles were unbiased (50% green and 50% purple). However, their locations were biased (with the same distributions as the other biased conditions). By decoupling the discriminated from the biased stimulus features (color and location, respectively) we were able to probe whether undiscriminated, low-level sensory location information alone biases subsequent location decisions, or whether making an intentional choice regarding the relevant feature is needed for serial-dependence to occur.

This condition included 38 participants, which consisted of two groups: (i) 15 of the participants who also performed Condition #3 (*Response Invariant*) and Condition #5 (*Alternative Feature*), and (ii) 23 naïve participants, to replicate the results of the former group. Results from two groups did not differ (*p* = 1.97, *t*(30) = 1.32, Cohen’s *d* = 0.475), and were therefore merged. Six participants were removed from this condition due to poor goodness-of-fit.

### Statistical analysis

Data analysis was performed with custom software using Matlab version R2014a (MathWorks) and the psignifit toolbox for Matlab, version 4 (Schütt et al., 2016). Psychometric plots were defined by the proportion of rightward choices as a function of circle location and calculated by fitting the data with a cumulative Gaussian distribution function. Separate psychometric functions were constructed for each prior type (center, left and right). The goodness-of-fit of the psychometric curves was evaluated using the Likelihood□ratio based *pseudo*□*R*□*squared* (Hosmer and Lemeshow, 1989; Menard, 2000), calculated by the proportional reduction in the deviance of the fitted psychometric model (*D*_*fitted*_) compared to that of the null model (*D*_*null*_):

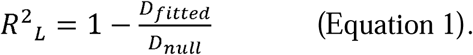

Deviance values for the fitted and null model (*D*_*fitted*_ and *D*_*null*_, respectively) were calculated using the psignifit toolbox. Only psychometric curves with *R*^*2*^_*L*_ > 0.5 were used for the final analysis. If any of a participant’s psychometric curves did not reach this level, that participant was removed from further analysis in that condition.

The point of subjective equality (PSE, i.e., the stimulus level with equal probability for making a rightward/leftward choice) was deduced from the mean (*μ*) of the fitted cumulative Gaussian distribution function. The effect of priors on subsequent location discriminations of test circles was assessed by calculating the difference in PSE between left and right prior types, as follows:

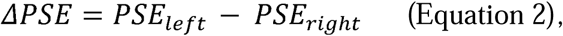

where the sign of ΔPSE corresponds the direction of the serial-dependence effect. Positive ΔPSE values indicate an attractive bias (i.e., subsequent perceptual choices are more likely to be the same as prior choices) and negative values indicate a repulsive (or adaptive) bias (i.e., subsequent perceptual choices are more likely to differ from prior choices).

Differences between PSEs of the different prior types (within a condition) was assessed using two-tailed paired t-tests, except for *Baseline A*, which included three types of priors (left, center and right) and was therefore assessed using a repeated measures one-way ANOVA with the Bonferroni correction for pairwise comparisons. ΔPSEs were compared between the different experimental conditions using one-way between-subjects ANOVA with pairwise comparisons adjusted with Bonferroni correction. Effect size was estimated by calculating Cohen’s *d* and η^2^ for *t*-test and ANOVA analyses, respectively.

### Model fits

While the ΔPSE (described above) depicts the aggregate behavioral effects of serial-dependence, it does not dissociate the effects of prior stimuli from prior choices. Therefore, to further dissect what elements of prior experience influence subsequent choices, we fit the data using a logistic regression model that separated the effects of prior stimuli from prior choices (Fig. 2). We fit the data from five out of the six conditions (it could not be done for Condition #4, *Response Inhibition* because no location discriminations for prior circles were reported).

**Figure 2.**
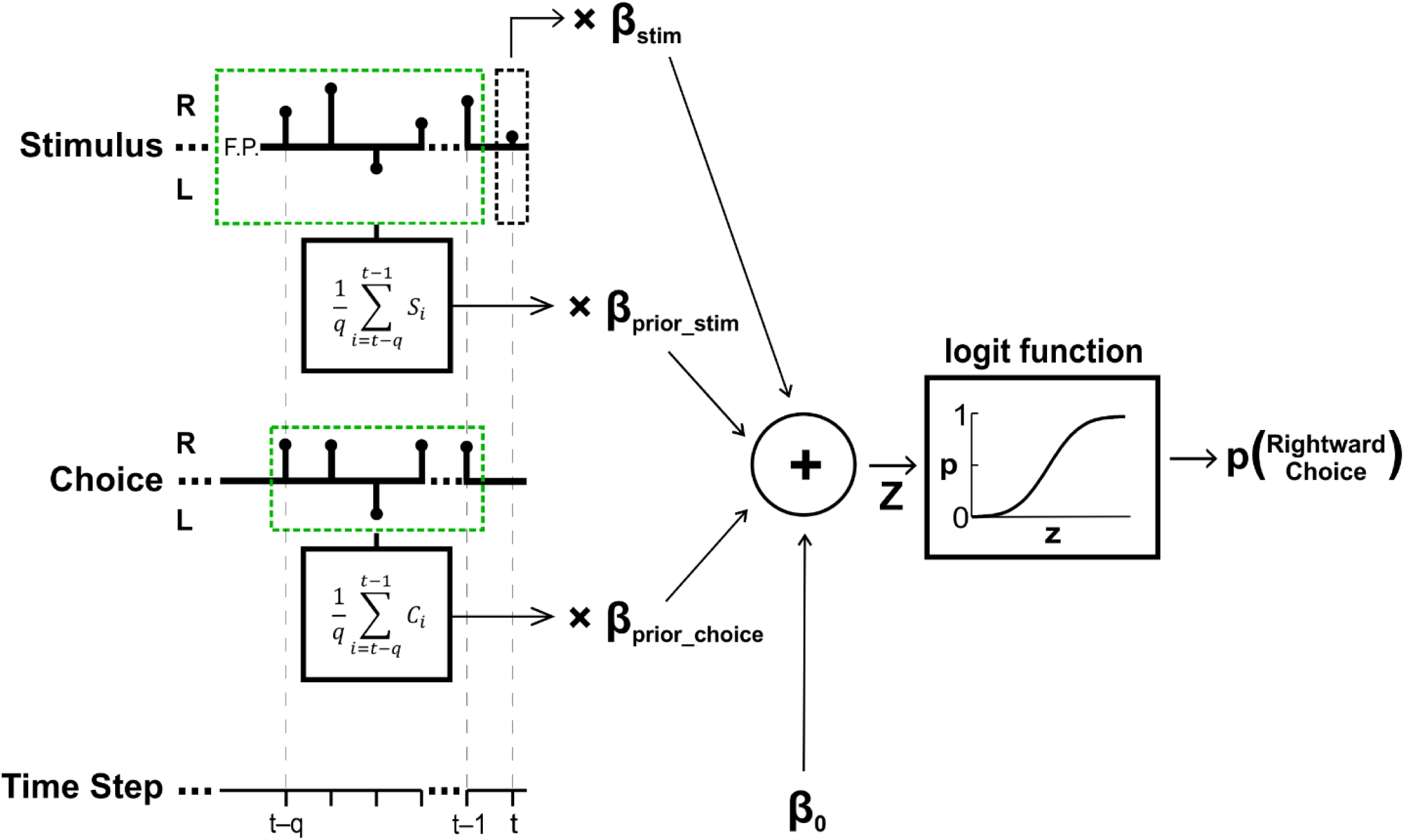
Perceptual decision model schematic. A logistic regression was used to model location discrimination choices based on four predictors (and their respective fitted beta coefficients): (1) current stimulus location (β_stim_), (2) stimulus location history (β_prior_stim_), (3) choice history (β_prior_choice_) and (4) baseline bias (β_0_). Black stems on the stimulus and choice axes mark stimulus (*S*_*i*_) and choice (*C*_*i*_) at time step *i* for an example trial, with test stimulus at *t* (black dashed box), and prior stimuli and choices *q* steps back (for example, priors biased to the right, green dashed boxes). The trial began with a central fixation-point (*F.P.*). The sum of the product of the predictors with their respective coefficients (*z*) is passed through a logistic function to yield the probability of making a rightward choice in response to the test stimulus.

The logistic regression model calculated the probability of making a rightward choice for the current circle stimulus, as follows:

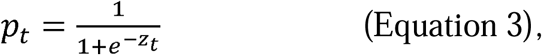

where *z*_*t*_ is a linear combination of different predictors that might affect the current perceptual decision (the specific *z*_*t*_ used for each model is described below). Choices were binary (right/left; coded by ‘1’ and ‘-1’, respectively), whereas circle locations (i.e., stimulus intensities) were graded along a continuum (with negative and positive values for leftward/rightward locations, respectively). To allow better comparison between the logistic regression weights of these two parameters, circle locations were normalized by the root-mean-square (RMS) of the actual location values presented in the experiment (separately for each experimental condition) such that both parameters had RMS = 1.

We tested and compared four competing models (separately for each experimental condition), that were based on the same logistic regression and differed only by their input parameters:

### M_1_: “No-History” Model

This model does not include information from previous trials and was used as a baseline for assessing the added value of incorporating prior information (in the following models). The linear combination of predictors used for M_1_ was:

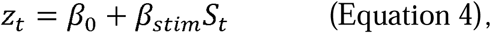

where *β*_0_ represents the observer’s individual baseline bias towards one of the two options (left or right), *S*_*t*_ is the location of the current stimulus and *β*_*stim*_ is its fitted weight.

### M_2_: “Stimulus-History” Model

M_2_ is like M_1_, but with the addition of prior stimulus information as a predictor:

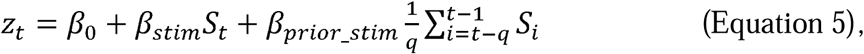

where 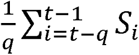 S is the averaged location of the *q* prior stimuli and *β*_*prior_stim*_ is the weight of this parameter. Setting *q* = 1 (‘1-Back’ analysis) enabled us to investigate the effects of just the previous stimulus and setting *q* = *Q* (‘*Q*-Back’ analysis; where *Q* reflects the number of priors presented on a given trial) enabled us to investigate the aggregate effect of the prior stimuli. In this latter case, our aim was not to quantify the individual effects of each prior stimulus (as is sometimes done, at the cost of additional parameters), but rather to quantify their average affect. This takes advantage of our paradigm design, with consecutively biased priors (interleaved on a trial-by-trial basis), without the need to add more parameters.

### M_3_: “Choice-History” Model

M_3_ is like M_1_, but with the addition of prior choice (not prior stimulus) information as a predictor:

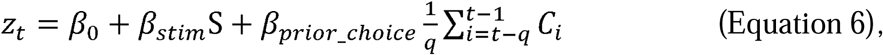

where 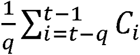 is the rightward choice ratio for *q* priors and *β*_*prior_choice*_ is the weight of this parameter. As before, *q* = 1 was used to investigate the effects of just the previous choice and *q* = *Q* was used to assess the aggregate effect of the prior choices.

### M_4_: “Stimulus and Choice History” Model

Finally, in order to estimate the specific effects of prior stimuli and prior choices on subsequent perceptual decisions, M_4_ included both prior stimuli and prior choices (in addition to the baseline parameters of M_1_):

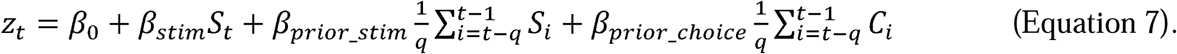

Here too, *q* = 1 was used to investigate the effects of just the previous prior and *q* = *Q* was used to assess the aggregate effect of the priors.

### Model Comparison

We used the Bayesian Information Criterion (BIC) to compare the models. The BIC for a given model (*M*_*i*_) is defined by:

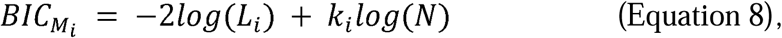

where *N* is the number of observations, in this case, the number of trials (or ‘test’ circles) within an experimental condition. The number of free parameters of *M*_*i*_ is represented by *K*_*i*_, and *L*_*i*_ is its maximum-likelihood. A lower BIC value represents a more likely model (Schwarz, 1978; Raftery, 1995). BIC values were calculated, per participant, for each of the models. To compare two given models, we calculated the difference between their BIC values (ΔBIC). ΔBIC magnitudes between 2 and 6, between 6 and 10 and > 10 are considered positive, strong and very strong evidence (respectively) for the model with the lower BIC value (Raftery, 1995; Wagenmakers, 2007; Jarosz and Wiley, 2014).

## RESULTS

Results from our location discrimination paradigm, designed to investigate what aspects of prior perceptual decisions lead to serial-dependence (six conditions, summarized in Table 1) provide a broad view of serial-dependence, and specifically implicate “task relevance” as a pivotal feature for the emergence of serial-dependence.

### Location discriminations are biased towards recent choice history

In the standard form of the paradigm (*Baseline*, Condition #1) the same task, using the same stimuli, was applied both to the prior and test discriminations. Specifically, participants were required to discriminate whether the stimulus (a solid white circle) lay to the right or to the left of the screen center (2AFC). Stimulus locations were biased to the right or to the left for the prior discriminations, followed by unbiased (i.e., balanced) test stimuli. Responses to these test stimuli were sorted by prior bias, analyzed and fit with psychometric curves.

Results from an example participant (Fig. 3A) expose substantial serial-dependence. Following rightward biased priors, test stimuli were more likely to be discriminated rightward (green curve). This is best discerned by the value of the psychometric function (> 0.5, i.e., greater likelihood for rightward choices) at the ambiguous stimulus x = 0° (where it crosses the vertical dotted line). This rightward choice bias is also depicted by a negative PSE – the green curve crosses the horizontal dashed line (y = 0.5) at a negative x-value. Similarly, leftward biased priors (orange curve) led to a leftward bias of subsequent test discriminations (and a positive PSE). Responses following unbiased priors (gray curve) lay in between the other two.

**Figure 3.**
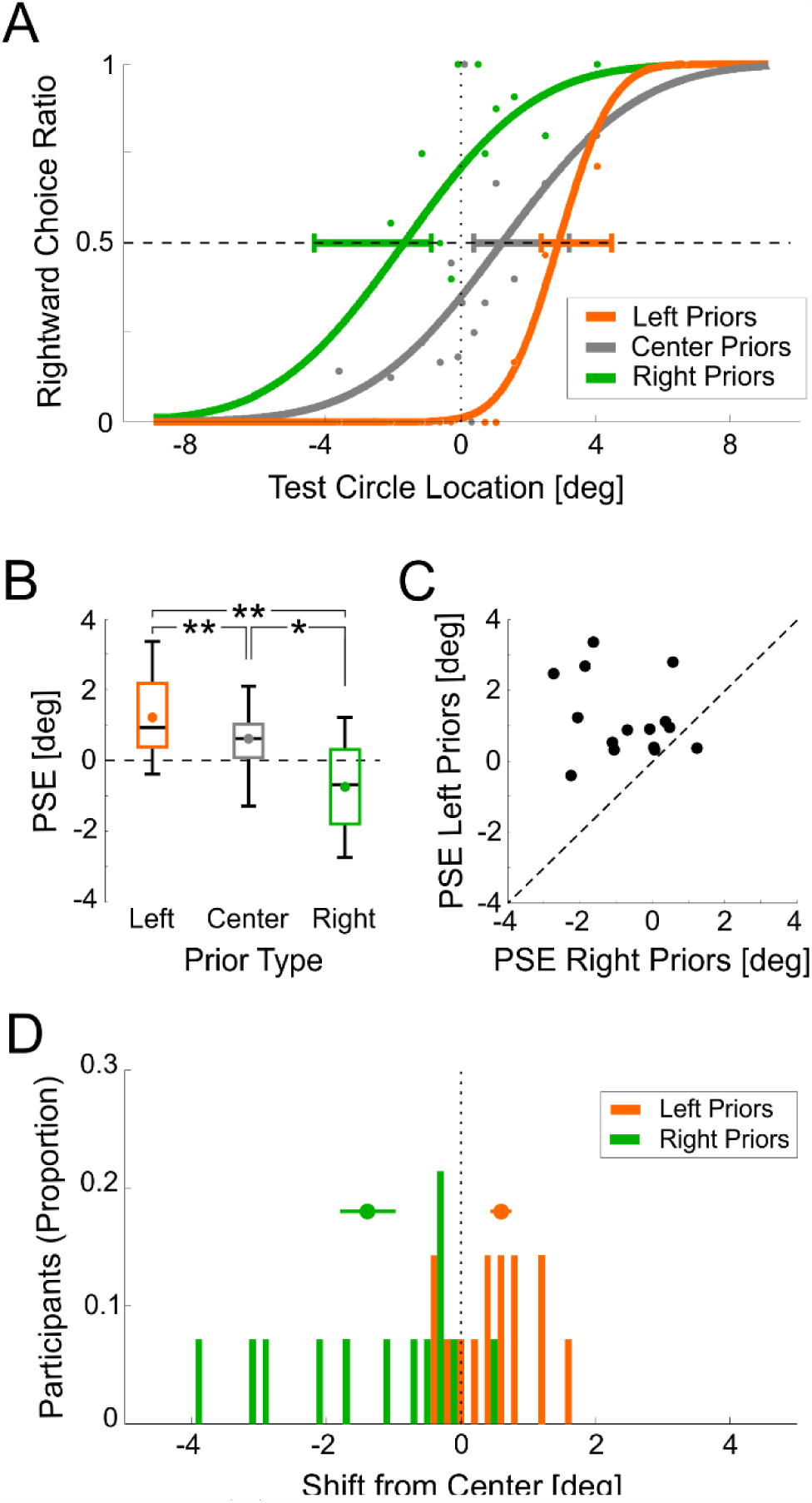
Baseline A condition results. (A) Psychometric functions are presented for an example participant, sorted by prior type: left, center and right (orange, gray and green, respectively). The data points represent the proportion of rightward choices (for the test circles only) fitted (per prior type) with a cumulative Gaussian distribution function (solid lines). The horizontal dashed line marks the point of subjective equality (PSE; y = 0.5), and the vertical dotted line marks the screen center (x = 0). Horizontal error bars mark 95% confidence intervals of the PSE. (B) Summarized PSEs of the group data for the three prior types: left, center and right (orange, gray and green, respectively). The median and range of the data are represented by the black lines and whiskers, respectively. The colored dots represent the respective means. (C) PSEs for left vs. right prior types. Each data point represents one participant. The diagonal line marks y=x. (D) Distribution of PSE shifts across participants, calculated by subtracting the unbiased (center) PSE from the left and right prior type PSEs (orange and green, respectively). The objective screen center (x=0) is represented by the vertical black dotted line. Above the histograms, mean ± SEM are presented by the circle markers and horizontal lines, respectively. *p<0.05, **p=0.05.

Robust serial-dependence was also seen across the group (Fig. 3B-D). Responses to the test stimuli were biased leftward (positive PSEs; orange box in Fig. 3B) following leftward biased priors, and biased rightward (negative PSEs; green box) following rightward biased priors. PSEs following unbiased priors (center, gray box) lay between the other two. PSEs differed significantly by prior type (within-subjects one-way ANOVA, *F*(2,28) = 15.03, *p* = 0.000037, η^2^ = 0.518). And, post-hoc Bonferroni pairwise comparisons revealed significant differences between all three prior types (*p* = 0.011 for the left vs. center, *p* = 0.038, for the right vs. center, and *p* = 0.003 for the right vs. left). A subject-by-subject comparison of PSEs demonstrates high consistency and robustness across participants – all (but one) had a larger PSE following left (vs. right) biased priors (14 out of 15 datapoints lie above the diagonal dashed line in Fig. 3C). Figure 3D presents the distribution of PSEs for leftward and rightward biased priors (orange and green, respectively) in relation to each individual’s unbiased/center PSE.

For better comparison with the other conditions (Conditions #2-5, described below) a second version of the *Baseline* (Condition #1, variant “B”) was run in addition to that presented above (variant “A”). Variant B had 2 methodological differences: i) the number of priors was random (between 1-9 vs. a constant 5 in variant A), and ii) only the two biased prior types (right and left) were tested (i.e., without the center/unbiased prior type, also tested in variant A). The results from variant B replicated those of variant A, and demonstrate similar and robust PSE shifts following biased priors. This was quantified by calculating the ΔPSE (the difference in PSE for leftward vs. rightward biased priors). Positive ΔPSE values indicate an “attractive” effect. Namely, priors biased leftward (rightward) increase the likelihood for subsequent leftward (rightward) choices (like the example in Fig. 3A). *Baseline* B variant, like *Baseline* A (see above), had a significantly positive ΔPSEs (Fig. 4A; *t*(13) = 3.61, *p* = 0.003, Cohen’s *d* = 0.966). Since variant B was better matched methodologically to the control conditions (#2-5), it was used for cross-condition comparisons (below).

**Figure 4.**
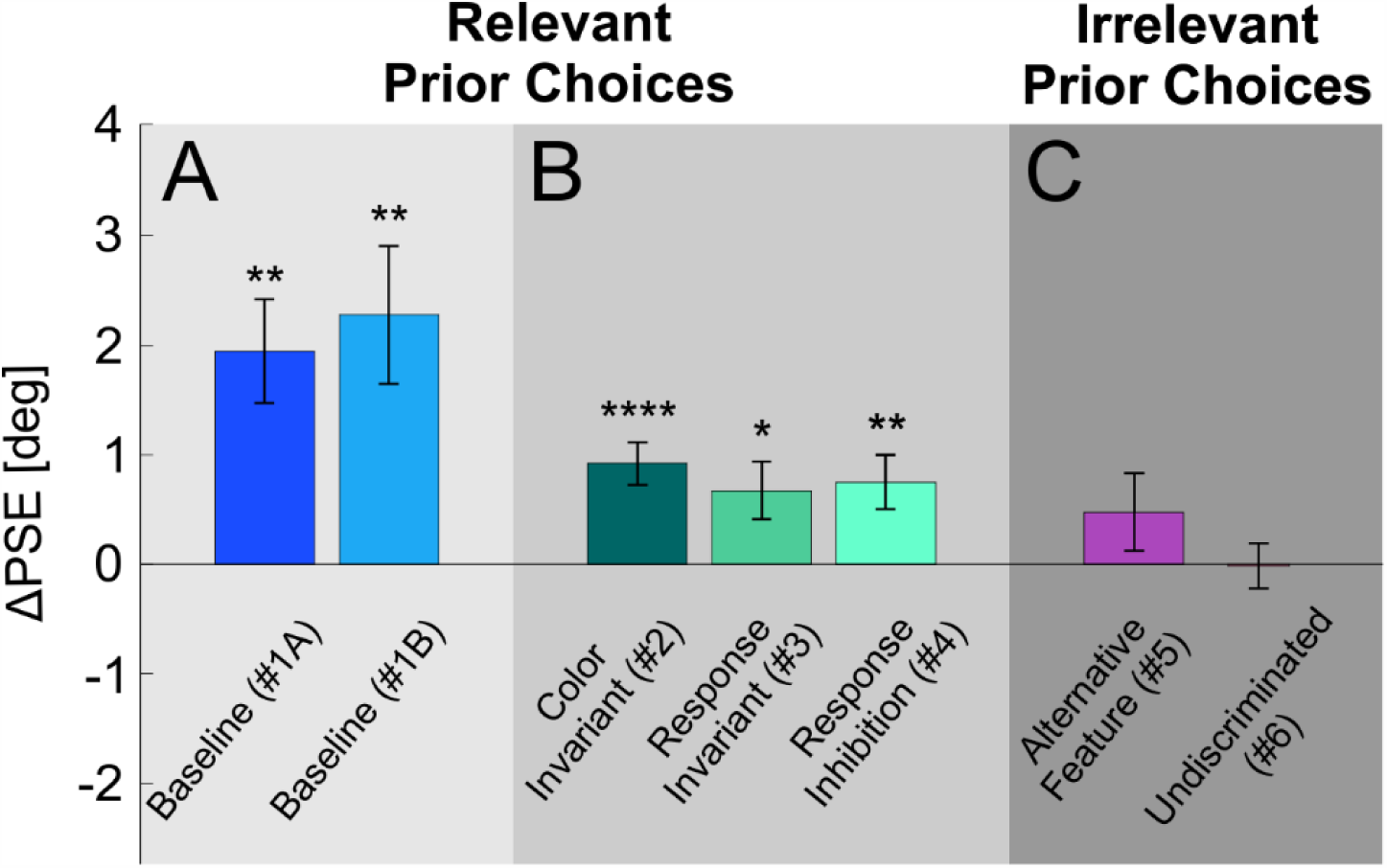
Priors effects across experimental conditions. The PSE difference between left and right prior types (ΔPSE = PSE_Left_priors_ – PSE_Right_priors_) is presented per condition (mean ± SEM). Condition names are presented below the bars (Condition number in parenthesis). Significant shifts are seen for: (A) the *Baseline* conditions (blue shades), and (B) control conditions in which the task for the prior stimuli was relevant to the test task (green shades). But, not for (C) conditions in which the task for the prior stimuli was irrelevant to the test task (pink shades). *p<0.05,**p<0.01,****p<0.0001.

### Relevance is key for serial-dependence

To test the effects of task relevance on serial-dependence, we developed five more (control) conditions based on the standard (*Baseline*) location discrimination task. These are divided into two main groups: the *relevant* control conditions (Conditions #2-4) and the *irrelevant* control conditions (Conditions #5-6). In the *relevant* control conditions, perceptual discriminations for both the prior and test stimuli were about the same feature (location). Other (irrelevant) features were manipulated to differ between the priors and test. Hence, we use the term “relevant” here to indicate that the same feature was discriminated for both prior and test stimuli. For the *irrelevant* control conditions, a different feature was discriminated for the prior stimuli (color) vs. the test stimuli (location). These conditions allowed us to test the effects of other biases, specifically, motor selection biases, and low-level sensory biases (when location was not discriminated).

By the term “relevant”, we don’t imply that there was indeed any objectively relevant information in the priors that could be useful for discriminating the test stimulus locations (these were independent by design). Rather, we propose that serial-dependence likely reflects an implicit presumption of the subjects that statistics from previous trials can help subsequent discriminations. While this might not improve performance in most lab experiments (which generally present independent stimuli across trials) it might help in the real world, where statistical structure is not independent. Hence, we hypothesize here that serial-dependence emerges when prior and test discriminations are about the same (relevant) feature, but that irrelevant discriminations do not lead to serial-dependence.

Indeed, in all the three *relevant* control conditions significant serial-dependence was observed (Fig. 4B): In the *Color Invariant* condition (Condition #2) prior and test circles differed in terms of color. But, color was an irrelevant parameter, since participants were required to discriminate location for all stimuli. Significant PSE shifts were still observed (*t*(23) = 4.7, *p* = 0.000096, Cohen’s *d* = 0.961; Fig. 4B left bar). In the *Response Invariant* condition (Condition #3) discriminations for the prior and test circles were reported using different motor responses (up/down or right/left keys, respectively). This is a particularly interesting condition, since high-level location discriminations link the prior and test stimuli (stimulus-action mapping was different for the two). Nonetheless, significant PSE shifts were still observed (*t*(21) = 2.52, *p* = 0.019, Cohen’s *d* = 0.54; Fig. 4B middle bar). Finally, in the *Response Inhibition* condition (Condition #4) discrimination responses were reported only for the test circles, but withheld for the prior circles. Here too, significant PSE shifts were still observed (*t*(16) = 3.004, *p* = 0.008, Cohen’s *d* = 0.73; Fig. 4B right bar).

Hence, in all conditions where the priors’ task was the same as the test task (location discrimination; Conditions #1-4), serial-dependence was observed, even when other features were changed. Notably, this effect persisted even when current and prior choices were not reported or were reported by different actions. We will now see that this was only the case as long as the prior decisions themselves remained relevant.

### Irrelevant biases do not lead to substantial serial-dependence

In the *irrelevant* conditions, color was discriminated for the prior stimuli, while location was discriminated for the test stimuli. These conditions allowed us to examine the effects of other types of prior biases. Firstly, the effect of motor repetition was tested by generating choice biases for rightward or leftward choices in the priors’ (color discrimination) task (Condition #5 *Alternative Feature*). Colors were reported using the same left/right arrow keys as the subsequent location choices for the test stimuli, thereby biasing prior motor responses. Motor biasing led to a small (but not significant) positive serial-dependence effect. (*t*(19) =1.33, *p* = 0.199, Cohen’s *d* = 0.29; Fig. 4C left bar).

In the last, *Undiscriminated* condition (Condition #6), the prior circles locations were biased, but participants did not discriminate location for the prior circles. Rather, they discriminated color. Hence the prior location biases are considered low-level and undiscriminated. In this condition as well, no significant PSE shift was seen (*t*(31) = −0.1, *p* = 0.92, Cohen’s *d* = −0.018; Fig. 4C right bar). Notably, the *Undiscriminated* condition (Condition #6) is identical to the *Color Invariant* condition (Condition #2) in every aspect, except for one: the discriminated feature of the prior stimuli (color for the former, and location for the latter). Yet, significant serial-dependence is only seen for the *Color Invariant* (task relevant), but not the *Undiscriminated* (task irrelevant) condition. Thus, when prior decisions were irrelevant, there was no substantial serial effect, even when motor actions or lower-level sensory characteristics of the prior trials were biased.

### Similarity boosts serial-dependence

We have shown above that task relevance has a key role in biasing sequential perceptual decisions. We now examine whether the degree of similarity (even of irrelevant features) affects the magnitude of serial-dependence. Greater similarity between two events increases our belief that they originate from the same distribution (Tversky and Kahneman, 1973). Thus, even though we have shown that on their own, irrelevant features do not elicit serial-dependence, greater similarity between events, even if irrelevant, might increase the effect of serial-dependence.

Of the four *relevant* conditions, the *Baseline* condition (Condition #1) had the greatest similarity between prior and test discriminations (these were essentially identical from the point of view of the participant, who simply had to discriminate location for a series of white circles). All the other *relevant* conditions (Conditions #2-4) had specific dissimilarities between prior and test discriminations. Comparing ΔPSEs across these conditions (#1-4; Fig. 4) indeed revealed significant differences (between subjects one-way ANOVA, *F*(3,73) = 4.478, *p* = 0 .006, *η*^*2*^ = 0.155). Using post-hoc pairwise comparisons with Bonferroni adjustment we now assess the effects of specific task dissimilarities.

First, the contribution of stimulus similarity was assessed by comparing the *Baseline B* Condition #1 and *Color Invariant* Conditions #2. While the prior and test stimuli were identical in Condition #1 (white circles), they differed from one another in Condition #2 (green or purple for the prior stimuli, and white for the test stimuli). All other parameters were the same. The *Baseline* Condition #1 had significantly larger ΔPSEs vs. the *Color Invariant* Condition #2 (*p* = 0.03; post-hoc pairwise comparison with Bonferroni adjustment). Therefore, although identical stimuli are not necessary to trigger serial-dependence, stimulus similarity does boost the effect.

The contribution of response similarity was assessed by studying the effects of two *relevant* conditions in which responses were manipulated: the *Response Invariant* Condition #3 (in which discriminations were reported using different responses) and the *Response Inhibition* Condition #4 (in which responses were withheld for prior stimuli). To assess the effects of reporting discriminations using different responses, Condition #3 was compared to the *Color Invariant* Condition #2. These two conditions were otherwise identical (besides the requirement to use different keys in Condition #3). No significant difference was seen between the ΔPSEs of these two conditions (post-hoc pairwise with Bonferroni *p* > 1). Therefore, serial-dependence was still strong even when motor responses were dissociated. In a more nuanced analysis (below) that separates the effects of prior choices and stimuli we do find that using the same motor actions slightly enlarges the effects of serial-dependence (see section *Relevant choices bias subsequent discriminations*). This is in line with the trend that we observed for a small motor repetition effect. Thus, motor similarity seems to magnify, (but not to be required for serial-dependence.

To assess the effects of withholding responses, *Response Inhibition* Condition #4 was compared to the *Baseline B* Condition #1. These two conditions differed by whether/not responses were reported for the prior stimuli. In Condition #4, participants were instructed to only respond to the last stimulus. Because the number of priors was variable, participants had to discriminate all stimuli, but only reported their discrimination for the last (test) stimulus. Other parameters were the same (stimuli were all white). The *Baseline* Condition #1 had significantly larger ΔPSEs vs. Condition #4 (*p* = 0.02, post-hoc pairwise comparison with Bonferroni adjustment). Although the same participants took part in both conditions, for consistency with the other comparisons in this section a between-subjects comparison was used (a within-subjects comparison provided similar results: *p* = 0.016). Therefore, actively reporting prior discriminations boosts serial-dependence.

### Choices bias subsequent discriminations

Quantifying shifts in the psychometric curves (ΔPSE), as presented above, is a useful tool to expose serial-dependence. Nonetheless, aggregating the effects into a single overall psychometric shift, has limited capacity to answer deeper questions. Specifically, what aspect of prior discriminations drives the phenomenon – prior choices, prior stimuli, or a combination of the two? To answer this question, we fit the data on a trial-by-trial basis with a logistic-regression model, that could account separately for the effects of prior choices and prior stimuli (see Fig. 2 and Methods section *Model fits* for details). For comparison, we fit the logistic regression with four different combinations of parameters. All four had two standard parameters (to account for the effects of the current stimulus and a baseline bias), and differed by whether or not they also took into account prior choices and/or prior stimuli, as follows: (M_1_) no history (no prior choice/stimulus information), (M_2_) stimulus history only, (M_3_) choice history only or (M_4_) both stimulus and choice history. Models fits were performed per participant, and condition (except for Condition #4, which was not fit because choices were not reported for prior stimuli).

To quantify the value of including prior information, we calculated the difference in Bayesian Information Criterion (ΔBIC) between each of the history models (M_2-4_) vs. M_1_ the no-history model (Fig. 5; diagonal-stripe, filled and vertical-stripe bars, for M_2_, M_3_ and M_4_, respectively). For the *relevant* conditions (#1-3; Fig. 5A and B) ΔBICs are large and negative, indicating better model fits with history (the vertical black dashed line at ΔBIC = −10 marks very strong evidence for the models with history; Raftery, 1995; Wagenmakers, 2007; Jarosz and Wiley, 2014).

**Figure 5.**
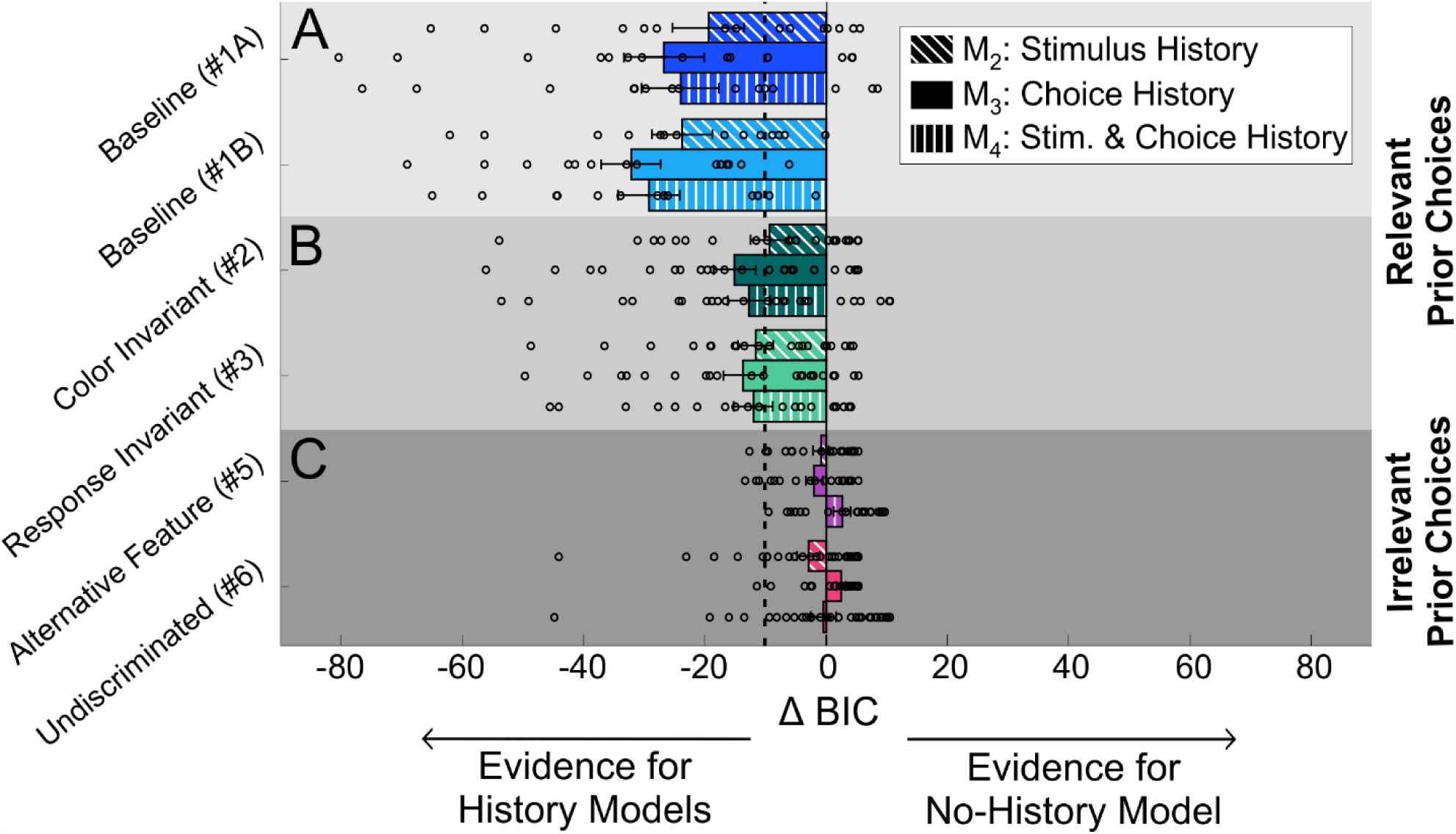
Model Comparisons. The Bayesian information criterion (BIC) is presented for the three history models in relation to the ‘no history’ model, across experimental conditions. ‘Stimulus history’, ‘choice history’ and ‘stimulus and choice history’ models are represented by the diagonal-stripe, filled and vertical-stripe textured bars (ordered: top, middle and bottom within each condition), respectively. Negative ΔBIC values (ΔBIC = BIC_M*i*_ - BIC_M1_; where M_1_ represents the ‘no-history’ model and M_*i*_ represents one of the ‘history’ models, *i* = 2, 3 or 4) indicate more evidence for the ‘history’ model. ΔBIC < −10 is considered very strong evidence for the history model (the vertical dotted line marks ΔBIC = - 10). Circles represent ΔBIC values of individual partcipants, and bars and error bars mark the mean ± SEM. Choices were better explained by models with history for: (A) the baseline conditions and (B) control conditions in which the task for the prior stimuli was relevant to the test task, but not for (C) conditions in which the task for the prior stimuli was irrelevant to the test task.

By contrast, for the *irrelevant* conditions (#5-6; Fig. 5C), ΔBIC values are small, indicating no clear preference for the models with history. Namely, history information in the *irrelevant* conditions does not improve model performance. The results presented here used ‘*Q*-back’ model fits (i.e., all the priors in a trial). Using ‘1-Back’ fits (including only the most recent prior) also showed a preference for the history models in the relevant conditions, but with smaller ΔBIC values (ΔBIC < 8 compared to the large ΔBIC values seen in Fig. 5). Hence, we present results of the ‘*Q*-Back’ model fits, which take advantage of our paradigm design. We address this further below, in section *Accumulation effect of similar discriminations*.

Comparing between the history models in the *relevant* conditions (#1-3; Fig. 5A and B) shows better performance for M_3_ (choice history only) vs. M_2_ (stimulus history only) or even M_4_ (both stimulus and choice history). This suggests that prior choice is a more important factor than prior stimulus. To further expose this, we compared the beta coefficients of prior stimulus vs. prior choice from M_4_, which fit both parameters simultaneously (these could be compared because the stimulus and choice parameters were normalized by RMS before model fitting). Prior choices had a larger influence vs. prior stimuli (Fig. 6A). The data (and crosses, which represent the coefficients’ means ± SEM) lie off to the right of the y-axis (indicating large prior choice effects), but not far from the x-axis (indicating minor effects of prior stimuli). This is especially evident in the *relevant* conditions (#1-3; blue and turquoise shades) for which the prior choice beta coefficients were significantly positive (one sample *t*-test, *t*(14) = 9.75, *t*(13) = 6.91, *t*(23) = 7.93 and *t*(21) = 6.88, for Conditions #1A, 1B, 2 and 3, respectively, *p* < 0.00003 for all, after Bonferroni correction for multiple comparisons). By contrast, their prior stimulus beta coefficients were small (negative) and not statistically significant (*t*(14) = −0.8, *t*(13) = −0.8, *t*(23) = −2.73 and *t*(21) = −0.66, for Conditions #1A, 1B, 2 and 3, respectively, *p* > 0.28 for all, after Bonferroni correction for multiple comparisons).

**Figure 6.**
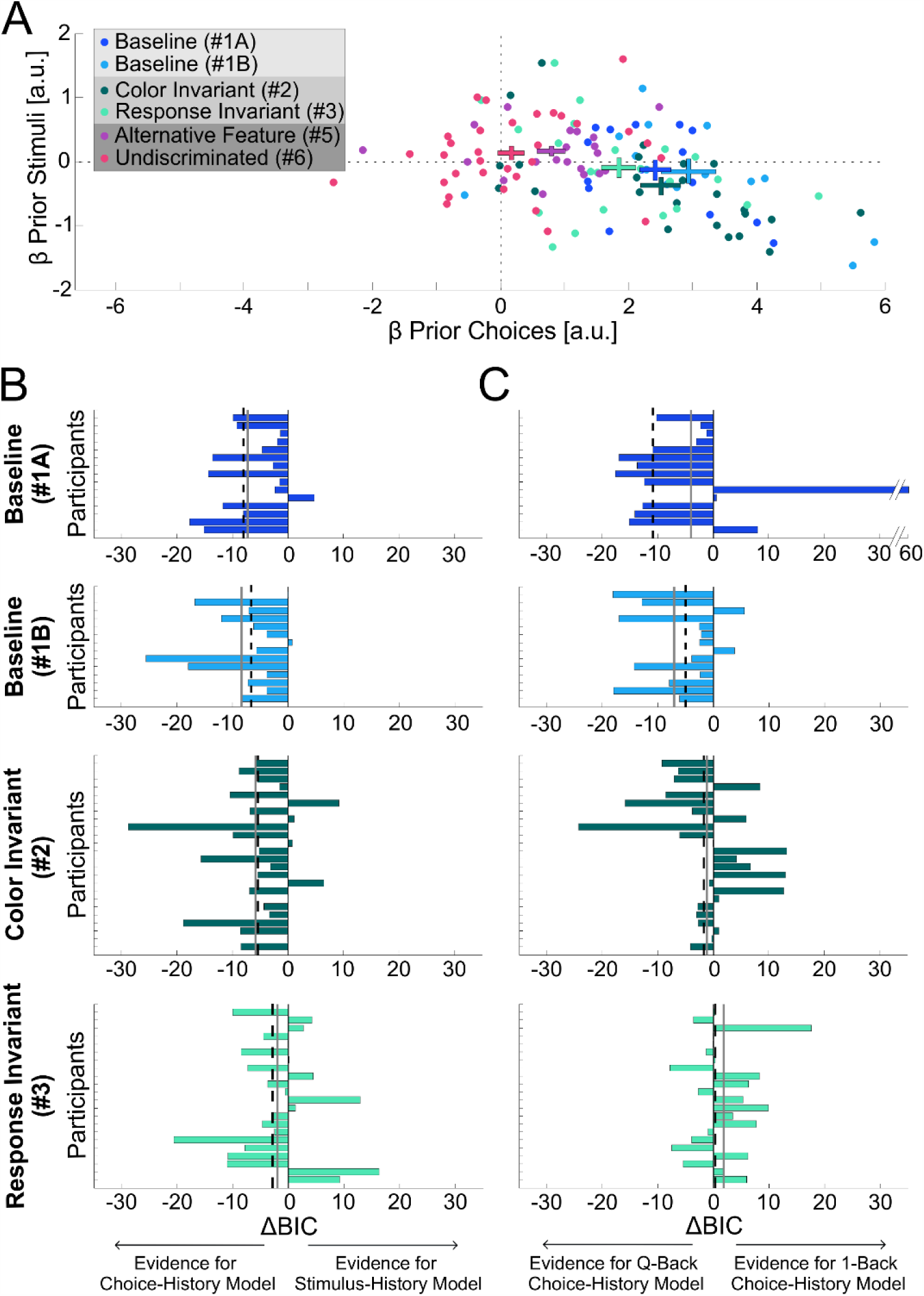
Choice vs. stimulus history comparison. (A) Beta coefficients for the prior stimulus vs. prior choice (from the ‘stimulus and choice’ history model M_4_) are plotted for each participant (data points), per condition (color coded). Crosses represent the mean ± SEM per condition. (B) Comparision between ‘choice-history’ (M_3_) and ‘stimulus-history’ (M_2_) models for each participant in the four ‘relevant’ conditions. Negative ΔBIC values mean more evidence for the ‘choice-history’ (vs. ‘stimulus-history’) model. Each bar represents the ΔBIC value for a single participant.The gray solid and black dashed lines represent the mean and median, respectively. (C) Comparision between ‘Q-Back’ and ‘1-back’ choice-history models for the four ‘relevant’ conditions. Negative ΔBIC values mean more evidence for the ‘Q-Back’ (vs. ‘1-Back’) model.

Comparing prior choice coefficients across all conditions revealed a significant difference (between subjects one-way ANOVA, *F*(5,121) = 16.595, *p* = 0.0000001, *η*^*2*^ = 0.407). Specifically, they were significantly smaller in the *irrelevant* vs. the *relevant* conditions. Post-hoc pairwise comparisons between each of the *irrelevant* conditions (Conditions #5 and #6) vs. each of the *relevant* conditions (Conditions #1-3) all revealed significant differences (Fig. 6A; *p* < 0.004 for all, with Bonferroni adjustment). Although smaller (compared to the *relevant* conditions) prior choice coefficients were nonetheless still significantly positive for Condition #5 (purple cross; *t*(19) = 3.54, *p* = 0.048 after Bonferroni correction for multiple comparisons). However, they were small (and not statistically significant) for Condition #6 (pink cross; one sample *t*-test, *t*(31) = 0.84, *p* = 0.4, not Bonferroni corrected). Prior stimulus coefficients were small and not significant also for the *irrelevant* conditions (one sample *t*-test; *t*(19) = 1.95, *p* = 0.065 and *t*(31) = 1.23, *p* = 0.23 without Bonferroni correction, for Conditions #5 and #6, respectively).

Calculating ΔBIC between the choice history model (M_3_) and the stimulus history model (M_2_), on a subject-by-subject basis, replicates the observation that prior choices affect subsequent decisions more than prior stimuli. *Baseline* Conditions #1 A and B have large, negative ΔBIC values on average, (negative for 90% of participants; Fig. 6B, top two rows). However, this seems to become less pronounced as the prior and test discriminations become less similar. Namely, in Condition #2 (where prior and test stimuli differed in color) ΔBIC values were still negative on average, but slightly less so (negative for 75% of participants; Fig. 6B, third row). And, in Condition #3, where prior and test stimuli differed in both color and motor responses, ΔBIC values were less negative (negative for 54.5% of participants; Fig. 6B, bottom row).

### Accumulation effect of similar discriminations

We have shown above that relevant prior choices elicit strong serial-dependence, and that greater similarity between the prior and subsequent events magnifies this effect. We now ask whether this effect is enhanced by a series of prior events or whether it is dominated by the most recent one. To address this, we examined whether the observers’ responses to test stimuli are better predicted by the aggregate effect of the *Q* prior choices in the current trial (‘*Q*-Back’) or by just the most recent one (‘1-Back’). We do not compare a model that fits each of the priors separately (due to the large increase in parameters this would entail; *Q*-Back and 1-Back have the same number of parameters). If *Q*-back provides a better fit, then this indicates an accumulation effect. A better 1-Back fit would suggest dominance of the most recent prior. This was quantified by calculating the ΔBIC between *Q*-Back and 1-Back model fits. The large negative ΔBIC values for the *Baseline* Condition #1 (Fig. 6C; top two rows) show a *Q*-Back preference, reflecting an accumulation effect of prior choices (negative ΔBIC values on average, and for 80% of participants). Here too, this effect decreased as similarity of the priors decreased. Namely, the ΔBIC values were on average closer to zero for Conditions #2 and #3 (Fig. 6C, third and fourth row, respectively) no longer showing a preference for *Q*-Back.

An accumulation effect of priors might suggest that a larger number of priors should lead to an increased serial-dependence effect. To test this hypothesis, we divided the trials into two groups – those with *less* (*Q* < 5, i.e.: 1-4) or *more* (*Q* ≥ 5; i.e.: 5-9) priors, and compared their ΔPSEs, for each of the *relevant* conditions (#1-4). Moreover, based on the *Q*-back analysis (above) this should be stronger for Condition #1. Indeed, we found that the difference between *less* or *more* priors was dependent on condition (*F*(3,73) = 2.863, *p* = 0.043, *η*^*2*^ = 0.102, two-way ANOVA interaction). Specifically, only for Condition #1, trials with *more* priors elicited stronger serial-dependence (ΔPSE mean ± SD = 2.12 ± 0.33°) compared to trials with *less* priors (1.19 ± 0.32°; *p* = 0.005; post-hoc pairwise comparison with Bonferroni adjustment). No significant differences were found in the rest of the conditions (#2-4, *p* > 0.28 for all; post-hoc pairwise comparisons with Bonferroni adjustment). These results echo those from the previous paragraph, and indicate that greater similarity of prior events leads to more accumulation of serial-dependence effects.

### Fixation-point reduces carry-over effects

Each trial began with a fixation point in order to reduce any carry over effects from the previous trial. To validate this assumption, we tested whether discriminations of the first prior of a trial (after the fixation point) were biased by perceptual history from the preceding trial (before the fixation point). Responses to these ‘first prior’ stimuli were sorted according to the prior bias from the previous trial, analyzed and fit with psychometric curves (in the same manner presented above for the test stimuli). Although a test discrimination lay in between the biased priors and the first prior of the next trial, its effect is limited because the stimulus was close to zero (due to the staircase) and was not biased, on average. This analysis was performed on the data from Condition #1 (*Baseline* variant B) because these data had large serial-dependence effects with accumulation, and in this condition, prior and test circles were indistinguishable to the participants, and the number of priors was random (so the appearance of a fixation-point could not be anticipated).

We found no significant difference between PSEs when separating the data by the prior type (left/right) of the previous trial (*paired t*-test; *t*(13) = −1.73, *p* = 0.1, Cohen’s *d* = −0.46). This suggests that the fixation-point indeed reduced the serial effects from previous trials. Similarly, assessing the effects of the previous trial (priors and test) on discriminating the first prior of the next trial using the logistic regression model (M_4_) showed a reduced effect of prior choices compared to that described for the test discrimination (prior choice beta coefficients mean ± SD = 1.31 ± 1.59° vs. 2.94 ± 1.59°; *p* = 0.004, *t*(13) = −3.5, Cohen’s *d* = −0.94). However, these were nonetheless statistically significant (one-sample t-test, *t*(13) = 3.08, *p* = 0.036, after Bonferroni correction). Hence, the effects of prior choices were reduced by more than half, but not eliminated. Like above (for the test circles) stimulus history beta coefficients were not statistically significant (mean ± SD = −0.21 ± 0.84°, *t*(13) = −0.95, *p* = 0.36, without Bonferroni correction).

## DISCUSSION

In this study, we found strong serial-dependence between sequential perceptual decisions, only when they were relevant to one another – namely, when discriminating the same perceptual feature. This effect was seen even when prior and current stimuli differed in their sensory attributes (e.g., color) or in the way the decision was reported. In a complementary manner, prior perceptual decisions about a different feature, irrelevant to the current decision, did not lead to substantial serial-dependence. Additionally, prior stimuli did not bias subsequent decisions about a feature, when that feature was not discriminated in the prior trials.

Our findings are in line with a recent study that found no serial-dependence when prior discriminations were about the variance of a stimulus value and subsequent discriminations about its mean (Suárez-Pinilla et al., 2018), considering variance and mean to reflect different features of the stimulus. But, in that study the different discriminations were reported using different motor actions. Hence, our results support and extend those findings by showing no serial-dependence between irrelevant decisions, even when they are reported using the same motor actions. Furthermore, we found that relevant prior decisions led to serial-dependence, even when these were reported using different motor actions (Condition #3, *Response Invariant*), as well as when motor actions were completely withheld (Condition #4, *Response Inhibition*). These results suggest that brain processes that determine whether prior decisions are relevant or irrelevant for subsequent decisions, do this independently of the actions used to report decisions. Importantly, prior perceptual decisions that are relevant, but not reported (and thus unobservable by an experimenter) can still elicit serial-dependence.

Fornaciai and Park (2018a, 2018b) found serial-dependence in numerosity perception, even when participants were told to ignore the prior stimuli, which were not relevant to the subsequent task. Their conclusion that task-irrelevant stimuli do elicit serial-dependence might seem, at face value, to contradict our results (and those of Suárez-Pinilla et al., 2018). However, closer inspection reveals that the results are actually consistent – the difference lies in the definition of “relevant.” In the numerosity task, the prior stimuli were indeed objectively irrelevant, since they contributed no information to solving the subsequent task. However, this is the case for almost all psychophysics tasks, in which stimuli are independent over trials. Yet serial-dependence is still observed, even though participants are instructed to discriminate just the stimulus presented (with the obvious implication that prior information is not relevant).

Rather, we interpret ‘relevance’ here to reflect the subjective experience – i.e., when the same task is performed in series, the observer will behave *as if* the prior stimuli are statistically relevant (even if they are objectively not). This might seem like not a good strategy (for laboratory tasks). However, given that consecutive events in the world are generally not independent, this may reflect a valid evolutionary skill. Our finding in this study, is that this happens if the prior decision is about the same stimulus feature. In the numerosity experiments (Fornaciai and Park, 2018a, 2018b), the prior stimuli lay embedded within a sequence of the same stimuli that were repeatedly discriminated. Thus, it is probable that also the prior stimuli were discriminated to some degree (even if participants were instructed to ignore them). The authors’ subsequent finding that attention to the prior stimuli is necessary to elicit serial-dependence supports this view. Accordingly, those experiments may be more similar to our Condition #4 (*Response Inhibition*; where choices were not reported) for which we also saw serial-dependence.

Our model analyses exposed that perceptual decisions in our study were biased specifically by, and towards, prior choices, not prior (low-level) sensory information. This is in line with recent studies that implicate prior choices as the cause of serial-dependence (Kaneko and Sakai, 2015; Braun et al., 2018; Talluri et al., 2018). Furthermore, when stimulus locations were biased (but these were not discriminated; Condition #6 *Undiscriminated*) serial-dependence was not observed. Together, these results indicate that it is the prior choices themselves, and not the low-level sensory stimuli, that led to serial-dependence. However, we also found that the effect of prior choices was modulated by sensory information. Specifically, serial-dependence was reduced when the stimuli differed in color. These results suggest an interesting interaction between prior choices and prior stimuli for serial-dependence: high-level, relevant, prior choices generate serial-dependence, which is modulated by low-level stimulus similarity.

One caveat regarding this conclusion, is that our task was one of location discrimination. Accordingly, by construction, subsequent stimuli appeared at different locations. Therefore, we cannot exclude the possibility that serial-dependence can also be triggered by low-level stimuli when they appear at the same spatial location. Hence, serial-dependence may have additional, low-level triggers. Nonetheless, our results demonstrate that relevant prior choices themselves generate robust serial-dependence, and that low-level location information (which is not specifically discriminated) does not lead to serial-dependence. Accordingly, these results differ from response “priming” (Klotz and Neumann, 1999) in terms of the specific influence of prior choices (vs. stimuli) and the higher-level conditions (i.e., relevant vs. irrelevant decisions) under which the effect is seen.

A minor effect of prior motor responses was found, when participants’ motor responses alone were biased (Condition #5, *Alternative Feature*). However, this was small compared to the effect of relevant prior choices. More importantly, serial-dependence was observed even when choices were not reported, in line with previous findings (Fischer and Whitney, 2014; Manassi et al., 2018). But, choice reporting did lead to a stronger effect. This action-related boosting of serial-dependence does not seem to result from general processes activated by motor actions, such as enhanced post-perceptual memory (Bliss et al., 2017; Fritsche et al., 2017) or enhanced attention to events that are acted upon, because serial-dependence was smaller when different actions were used to report choices. Rather, this (together with the findings above regarding stimulus similarity) seems to indicate that greater similarity between prior and current decisions boosts serial-dependence. Accordingly, stimulus-choice and/or choice-action coupling boosts serial-dependence.

With regards to low-level (stimulus or motor) similarities, it is possible that similarity binds events through categorization, such that similar events are labeled under the same category (Tversky and Kahneman, 1973). Prior events from the same (or close) category would be more likely to share their statistics, making them more useful to predicting upcoming stimuli, and perhaps, more relevant to the current goal. In turn, such prior events may lead to a larger serial-dependence. Indeed, a study by Petzold and Haubensak (2004) has shown that when participants were instructed to relate to stimuli with different colors as belonging to different categories, a serial-dependence effect emerged only within category. But, when participants were explicitly asked to ignore stimulus color and relate to all stimuli as belonging to the same category, the serial-dependence effect emerged within all types of stimuli. These findings support the idea that low-level similarities may contribute to the presumed relevance of prior choices, via a high-level categorization mechanism.

Neuronally, attraction to prior choices might result from residual activity in neuronal populations that encode prior choice, and therefore increase the probability for making the same choice again (Berlemont and Nadal, 2019). This theoretical mechanism is in accordance with animal studies that show priors and choice history activity, in posterior parietal cortex (Rao et al., 2012; Hwang et al., 2017), which might mediate subsequent biases via the basal ganglia (Lauwereyns et al., 2002; Hwang et al., 2019). Furthermore, choice, action and stimulus signals are mixed in posterior parietal cortex (Zaidel et al., 2017). Yet, the nature of this integration is unknown. It is therefore possible that posterior parietal cortex offers the high-level substrate to assess prior choice relevance, in light of low-level stimulus and action conjunctions, and biases subsequent choices via the basal ganglia.

In conclusion, our findings highlight high-level choice relevance as a critical trigger for serial-dependence. We also show that low-level sensory and motor information contribute to (but do not cause) the effect. Therefore, we suggest that serial-dependence reflects an active mechanism in the brain, in which relevant prior choices are used to predict and bias ongoing perceptual decisions toward the most likely interpretation of new incoming sensory information.

## Acknowledgments

This work was supported by The Israeli Centers of Research Excellence (I-CORE) program (Center No. 51/11) to AZ and by the Israel Science Foundation (ISF) grant # 673/17 to MB.

## Author Contributions

H.F and S.B developed the study with A.Z. All authors contributed to the study design. Testing and data collection were performed by H.F and S.B. Model-fit analysis was conducted by H.F. under the supervision of A.Z. H.F and S.B drafted the manuscript, and M.B and A.Z provided critical revisions. All authors approved the final version of the manuscript submission.

